# Inherited *BRCA1* epimutation as a novel cause of breast and ovarian cancer

**DOI:** 10.1101/246934

**Authors:** D. Gareth R. Evans, Elke M. van Veen, Helen J. Byers, Andrew J. Wallace, Jamie M. Ellingford, Glenda Beaman, Javier Santoyo-Lopez, Timothy J. Aitman, Diana M. Eccles, Fiona I. Lalloo, Miriam J. Smith, William G. Newman

## Abstract

**Background:** Pathogenic variants in *BRCA1* or *BRCA2* are identified in ~20% of families with multiple individuals with early-onset breast/ovarian cancer. Extensive searches for additional highly penetrant genes or alternative mutational mechanisms altering *BRCA1/2* have not explained the missing heritability. For the first time, we report transgenerational epigenetic silencing of *BRCA1* due to promoter hypermethylation in two families with breast/ovarian cancer.

**Methods:** *BRCA1* promoter methylation of ten CpG dinucleotides in breast/ovarian cancer families without germline *BRCA1/2* pathogenic variants was assessed by pyrosequencing and clonal bisulfite sequencing. RNA and DNA sequencing of *BRCA1* from lymphocytes was undertaken to establish allelic expression and the presence of germline variants.

**Findings:** *BRCA1* promoter hypermethylation was identified in two of 49 families with multiple women affected with grade 3 breast/high grade serous ovarian cancer. Soma-wide *BRCA1* promoter hypermethylation was confirmed in blood, buccal mucosa and hair follicles. Methylation levels were ~50%, consistent with the silencing of one allele and confirmed by clonal bisulfite sequencing. RNA sequencing revealed allelic loss of *BRCA1* expression in both families and this segregated with a novel heterozygous variant c.-107A>T in the *BRCA1* 5’UTR.

**Interpretation:** Our results indicate a novel mechanism for familial breast/ovarian cancer, caused by epigenetic silencing of the *BRCA1* promoter, segregating with an *in cis* 5’UTR variant in two independent families. We propose that methylation analyses are indicated in all families affected by early onset breast/ovarian cancer without a *BRCA1/2* pathogenic variant.

**Funding:** Funded by Prevent Breast Cancer (GA 12-006 and GA 15-002) and the Manchester NIHR Biomedical Research Centre (IS-BRC-1215-20007).

## Research in context

### Evidence before this study

In only ~20% of families with multiple individuals with early-onset breast/ovarian cancer are pathogenic variants in *BRCA1* or *BRCA2* identified. Since the discovery of these genes, extensive searches for additional highly penetrant genes or alternative mutational mechanisms altering *BRCA1/2* have not fully explained the missing heritability.

Epigenetic alterations have been proposed as an alternative mechanism to explain the missing heritability. This epigenetic mechanism has previously been described in familial colorectal cancer, due to inherited promoter hypermethylation of *MLH1* and *MSH2*. Methylation of the *BRCA1* promoter has been described in women with breast cancer but without a strong family history, but at low levels (<20%), and inheritance of promoter methylation has never previously been reported. Therefore, we investigated if inherited promoter methylation of *BRCA1* may be an alternative mechanism causing familial breast/ovarian cancer.

### Added value of this study

This is the first study that describes transgenerational promoter hypermethylation of *BRCA1*, resulting in complete transcriptional silencing of one allele. This hypermethylated allele segregates with a novel upstream variant in the 5’UTR, namely c.-107A>T. This novel mechanism may explain the missing heritability in some high-risk breast/ovarian cancer families.

### Implications of all the available evidence

Current clinical genetic testing approaches do not detect epigenetic changes that may account for some cases of familial breast/ovarian cancer. Detection of pathogenic variants is important to determine optimum treatment and appropriate cancer surveillance for at risk relatives. Therefore, methylation analysis in high-risk breast/ ovarian cancer patients negative for pathogenic variants in *BRCA1/2* may be informative.

## Introduction

Breast cancer is the commonest form of cancer in women.^1^ Germline heterozygous pathogenic variants in *BRCA1* and *BRCA2* account for 2-3% of all cases^2^ and up to 15% of cases of epithelial ovarian cancer.^3^ In families with multiple affected individuals with early-onset disease these percentages increase substantially, with *BRCA1* and *BRCA2* variants explaining approximately 20% of familial breast cancer and a higher proportion of familial ovarian cancer cases.^4^

Over the past twenty years, there have been exhaustive efforts to identify other breast and ovarian cancer susceptibility genes. This missing heritability has been postulated to be due to other highly penetrant genes, including *TP53*; genes of modest effect, including *PALB2/ATM*; or polygenic risks due to the combination of multiple variants each of small effect size.^5^ However, no other genes have been identified that confer a high risk of both breast and ovarian cancer.

Genetic testing by DNA sequencing and copy number analysis for pathogenic exonic variants in *BRCA1* and *BRCA2* is highly sensitive (estimated to detect over 90% of pathogenic variants)^6,7^ and is now offered routinely to individuals at high familial risk of breast/ovarian cancer. Previous studies by our group using RNA sequencing in high-risk families have shown that deep intronic variants in *BRCA1* or *BRCA2* do not contribute significantly to this mutational spectrum.^7^ Detection of pathogenic variants is important to determine appropriate cancer surveillance for at risk relatives; to reassure relatives without the familial causative variant regarding their risk and remove the burden of unnecessary screening; and to inform treatment choice, especially for poly ADP ribose polymerase (PARP) inhibitors.^8^ Gene promoter epimutations have been proposed as an alternative mechanism for the transcriptional silencing of cancer-associated genes.^9^ Promoter hypermethylation has been identified associated with tumor suppressor genes, both in the germline and as a somatic (acquired) event in tumor tissue,^9^ and results in transcriptional silencing.

Promoter hypermethylation of *BRCA1* is present in the tumor tissue of approximately 10% of sporadic breast cancers,^10^,^11^ in breast tumors of women with *BRCA1* germline pathogenic variants,^12^ and is more common in triple-negative (estrogen receptor, progesterone receptor, and HER2) breast cancer.^13^ Constitutional methylation of the *BRCA1* promoter has been reported in individuals with breast cancer,^14^ but this has always been at low ‘mosaic’ levels (maximum 20%) and there has been no convincing evidence that this has been inherited from one generation to the next. In contrast, inherited promoter hypermethylation of *MLH1*^15^ and *MSH2*^16^ has been reported in familial colorectal cancer. In this study, we describe the first breast/ovarian cancer families with an inherited germline variant that results in transcriptional silencing of *BRCA1* through promoter hypermethylation (secondary epimutation). This new mutational mechanism for *BRCA1* has important implications for diagnostic testing of patients at high-risk of breast/ovarian cancer and for optimum treatment selection.^17^

## Methods

### Patients and family members

*BRCA1* promoter methylation screening was undertaken in lymphocyte derived DNA of 49 unrelated individuals from families affected by breast/ovarian cancer and a Manchester score >34 without a germline *BRCA1*/*BRCA2* pathogenic variant. A Manchester score calculates the likelihood of detecting a pathogenic variant in *BRCA1*/*2*.^7,18,19^ In our local population 158 of 220 (71·8%) families with a Manchester score of >34 have had pathogenic variants in *BRCA1/2* identified by conventional genetic testing of DNA sequencing and multiplex ligation-dependent probe analysis (MLPA).

Blood, buccal mucosa, tumor, and hair samples were collected (where possible) from affected and unaffected family members with breast/ovarian cancer when *BRCA1* promoter methylation was detected. Cancer diagnoses were confirmed from hospital records or through the North West (England) Cancer Intelligence Service, which has data on all patients with any malignancy from 1960 onwards. DNA was extracted from blood by Chemagen (Perkin Elmer), hair using QIAamp DNA investigator kit (Qiagen), buccal mucosa on the Qiagen EZ1 system, and tumor using the Cobas® DNA Sample Preparation Kit (Roche). The study was approved by the Central Manchester Research Ethics Committee (10/H1008/24) and (11/H1003/3) and written informed consent obtained from each participant.

### *BRCA1* promoter methylation assays

Genomic DNA was bisulfite converted with EZ DNA methylation kit (Zymo Research) to distinguish between methylated and unmethylated DNA. *BRCA1* promoter methylation was determined by pyrosequencing (Qiagen) across 10 CpG dinucleotides within the *BRCA1* promoter to quantify the methylation status in DNA derived from hair follicles, buccal mucosal cells, peripheral blood lymphocytes, and tumor (eMethods).

Clonal bisulfite sequencing on a minimum of 37 clones was performed to determine if the methylation pattern was allele specific (eMethods).

### RNA and DNA analysis

To measure *BRCA1* expression, whole blood was collected in PAXgene Blood RNA tubes (PreAnalytiX) and RNA was extracted. RNA was converted to cDNA by RT-PCR using High-Capacity RNA-to-cDNA™ Kit (Applied Biosystems). Five SNPs in *BRCA1* exon 11 (rs1799949, rs16940, rs799917, rs16941, and rs16942) were genotyped by Sanger sequencing to determine if there was a difference in allelic ratios between the RNA and DNA genotypes to determine if there was silencing of one allele (eMethods, eTable 1).

### Haplotype analysis

To determine relatedness between families identified with *BRCA1* promoter methylation, 12 *BRCA1* intragenic SNPs were genotyped by Sanger sequencing to determine ancestral haplotypes (eMethods).^20^ In addition, genotyping using Affymetrix Genome-Wide SNP6.0 arrays was undertaken according to the manufacturer’s protocol. Genotypes and copy number data were generated within the Affymetrix Genotyping Console (v.4.1.3.840) via the Birdseed V2 algorithm and SNP 6.0 CN/LOH algorithm, respectively.

### Whole genome sequencing

PCR-free paired end whole genome sequencing (TruSeq DNA PCR-Free, Illumina) was undertaken on a HiSeqX platform. Reads were aligned against the human assembly GRCh38 via Burrows-Wheeler Aligner (BWA v.0.6.2) and variant calling using Genome Analysis Toolkit (3.4-0-g7e26428). Annotation was performed using v89 of the Ensembl database and compared to variation identified in the gnomAD dataset (http://gnomad.broadinstitute.org)^21^ (eMethods).

## Results

To determine if promoter hypermethylation of *BRCA1* could result in familial breast/ovarian cancer, methylation assays were undertaken. *BRCA1* promoter hypermethylation was identified in two women from a screen of 49 unrelated individuals with familial breast/ovarian cancer (and a Manchester score of >34) in whom previous Sanger sequencing and MLPA of *BRCA1* and *BRCA2* coding exons had not identified a pathogenic single nucleotide or copy number variant. In individuals with a Manchester score >34 there is a >70% likelihood of detecting a *BRCA1/2* germline pathogenic variant.^7,18,19^ The promoter hypermethylation was detected in a woman (Family-1; II-4, Figure 1A) with breast cancer aged 39 years, and a poorly differentiated serous ovarian cancer at 48 years with a strong family history of breast cancer (Manchester score 43) and a woman (Family 2, III-2, Figure 1B) with bilateral grade 3 triple negative breast cancer at age 38 and 46 years (Manchester score 35). In the two women, pyrosequencing assays on lymphocyte derived DNA were consistent with *BRCA1* promoter hypermethylation across 10 CpG dinucleotides (Figure 2A) (average 43% and 41%, respectively) indicating that one allele was fully methylated (Table 1, Figure 2B, C and eFigure 1). This hypermethylation pattern was consistent in DNA extracted from buccal mucosa (54% and 69%) and hair follicles (38% and 43%) (eTable 2), representing endoderm and ectoderm derived tissues, respectively. Clonal bisulfite sequencing orthogonally confirmed the *BRCA1* promoter hypermethylation pattern in the two affected women (Figure 2D).

**Figure 1:**
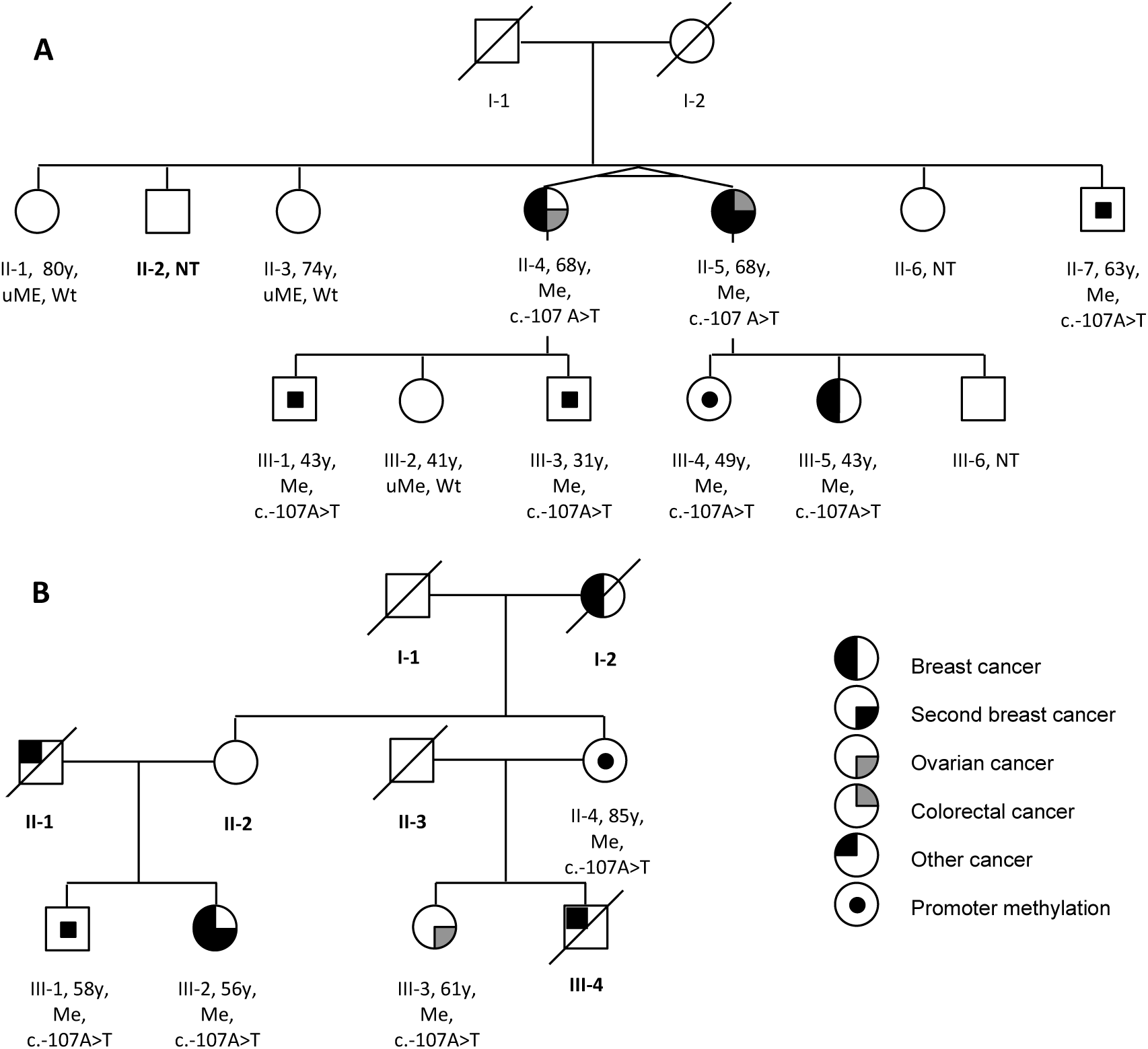
**A.** Pedigree of family 1. **B.** Pedigree of family 2. Y: years of age tested. uMe: unmethylated *BRCA1* promoter. Me: methylated *BRCA1* promoter. Wt: wild type.

**Figure 2:**
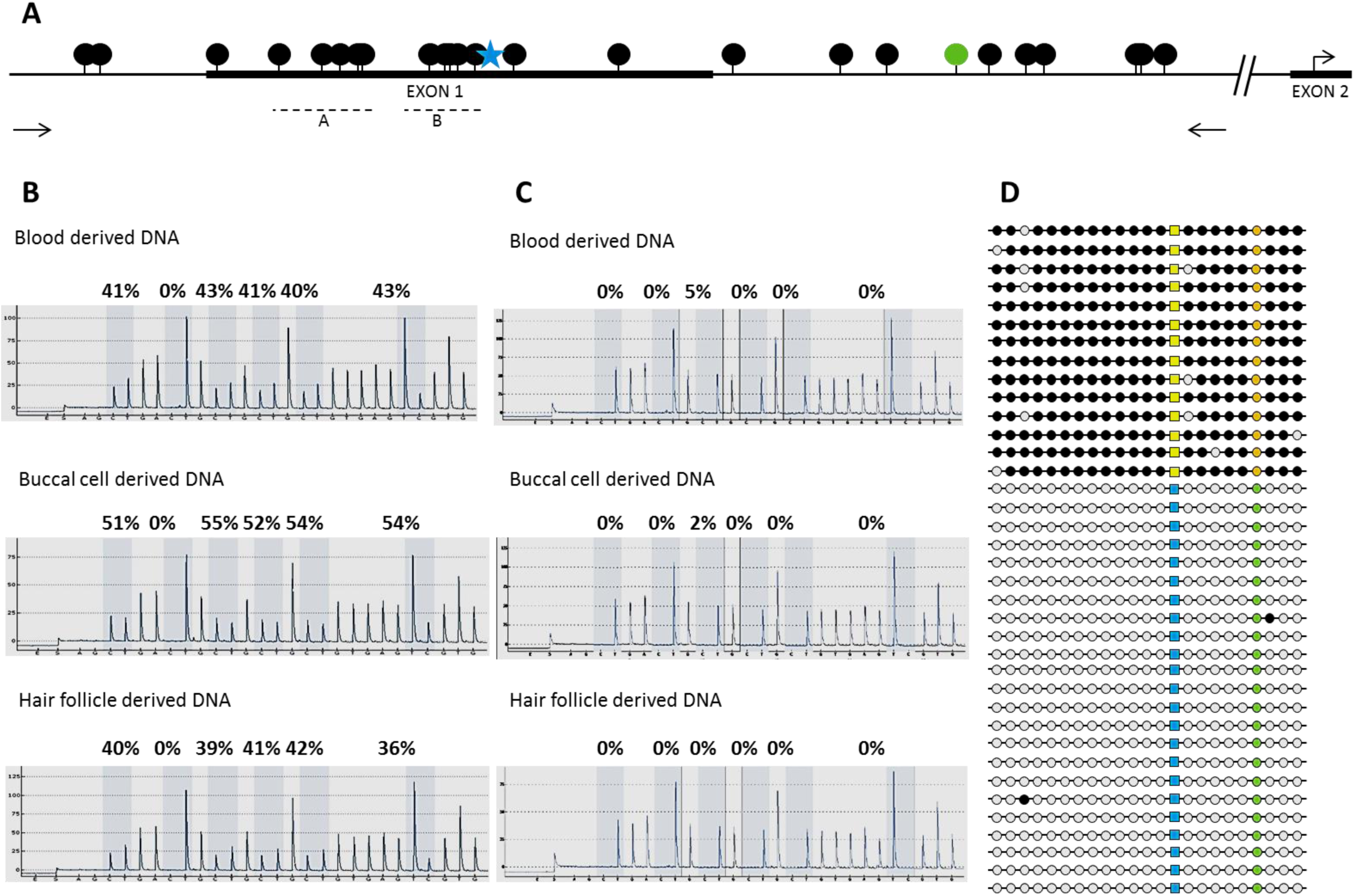
Schematic overview of *BRCA1* promoter region (black dots: CpG sites; star: c.-107; green dot: rs799905; arrows: primer locations for clonal bisulfite sequencing; dotted line: pyrosequencing region (A and B). **B and C.** Representative pyrograms (region B) show the level of *BRCA1* promoter methylation in lymphocytes, buccal mucosa and hair derived DNA of an affected and unaffected individual. Five CpGs and a control site (0%) (to ensure complete bisulfite conversion) are shaded and the percentage methylation as a ratio of C:T peak heights is calculated at each site (representing methylated versus unmethylated cytosine). **B.** Affected individual II-4 from Family 1. **C.** Unaffected individual II-1 from Family 1. Further pyogram data (region A) indicating methylation across the *BRCA1* promoter is available in the Supplementary appendix (eFigure 1). **D.** Schematic overview of clonal bisulfite sequencing results allelic discrimination is made based on rs799905 C>G (orange: G; green: C). The novel variant c.-107A>T is present on the methylated allele (yellow: T; blue: A; black is methylated, white is unmethylated).

**Table 1:** Summary of *BRCA1* promoter methylation status in lymphocyte derived DNA, clinical phenotype and genotype for the c.-107A>T variant for all tested individuals. M=male, F=female.

|  | *BRCA1* promoter methylation (mean %) | c.-107 A>T | Clinical Status (age at diagnosis years) | Sex | Age tested (years) |
| --- | --- | --- | --- | --- | --- |
| **Family 1** |  |  |  |  |  |
| II-1 | 1 | AA | Unaffected | F | 80 |
| II-3 | 0 | AA | Unaffected | F | 74 |
| II-4 | 43 | AT | Breast (39) and ovarian (48) cancer | F | 68 |
| II-5 | 37 | AT | Bilateral breast cancer (30 and 32), colorectal cancer (64) | F | 68 |
| II-7 | 41 | AT | Unaffected | M | 63 |
| III-1 | 38 | AT | Unaffected | M | 43 |
| III-2 | 1 | AA | Unaffected | F | 41 |
| III-3 | 44 | AT | Unaffected | M | 31 |
| III-4 | 41 | AT | Unaffected | F | 49 |
| III-5 | 32 | AT | Breast cancer (39) | F | 43 |
| **Family 2** |  |  |  |  |  |
| II-4 | 44 | AT | Unaffected | F | 85 |
| III-1 | 44 | AT | Unaffected | M | 58 |
| III-2 | 41 | AT | Bilateral breast cancer (38, 46) | F | 56 |
| III-3 | 43 | AT | Ovarian cancer (48) | F | 61 |

Segregation analysis for *BRCA1* promoter hypermethylation was undertaken in the two families. In Family 1 the proband’s identical twin (II-5), affected by bilateral grade 3 breast cancer at age 30 and 32 years (no receptor status available) and colorectal cancer at 64 years and II-5’s daughter (III-5), who had been affected by high-grade triple negative breast cancer at 39 years (Figure 1A) both had hypermethylation of the *BRCA1* promoter at similar allele frequencies to the proband (Table 1, eTable 2). Samples from the parents of the affected twins were not available, both were deceased, and neither had a history of cancer.

Samples from seven other family members were available (II-1, II-2, II-7, III-1, III-2, III-3 and III-4). Of these, four showed a soma-wide hypermethylated *BRCA1* promoter in blood, buccal mucosa, and hair follicles and in three a normal methylation pattern (Table 1, eTable 2, Figure 2B, C, and eFigure 1). In Family 2, the maternal first cousin (III-3) of the proband (III-2) had been diagnosed with high-grade serous ovarian cancer at 48 years, and also had soma-wide hypermethylation of the *BRCA1* promoter. The mother of III-2 was deceased (due to myocardial infarction at 76 years with no history of cancer). Her sister (II-4), the mother of III-3, was alive at 85 years, also with no history of cancer and had similar hypermethylation levels (43%) to her affected daughter and niece. The healthy brother (III-1) of the proband also showed hypermethylation of the *BRCA1* promoter.

DNA extracted from formalin fixed paraffin embedded breast tumor was available from individual III-5 (Family 1). Genotyping showed loss of the wild type allele across five informative intragenic SNPs (eTable 3) (i.e. only the alleles of the variants not expressed in the cDNA were present), consistent with loss of *BRCA1* as the second hit in the tumor tissue. Expression analysis of *BRCA1* in RNA extracted from lymphocytes was undertaken in individuals with promoter hypermethylation. Absence of heterozygosity across five SNPs with high minor allele frequencies within the *BRCA1* cDNA suggested allelic imbalance (Figure 3A), secondary to loss of expression of one allele due to hypermethylation of the *BRCA1* promoter (Figure 3C).

**Figure 3:**
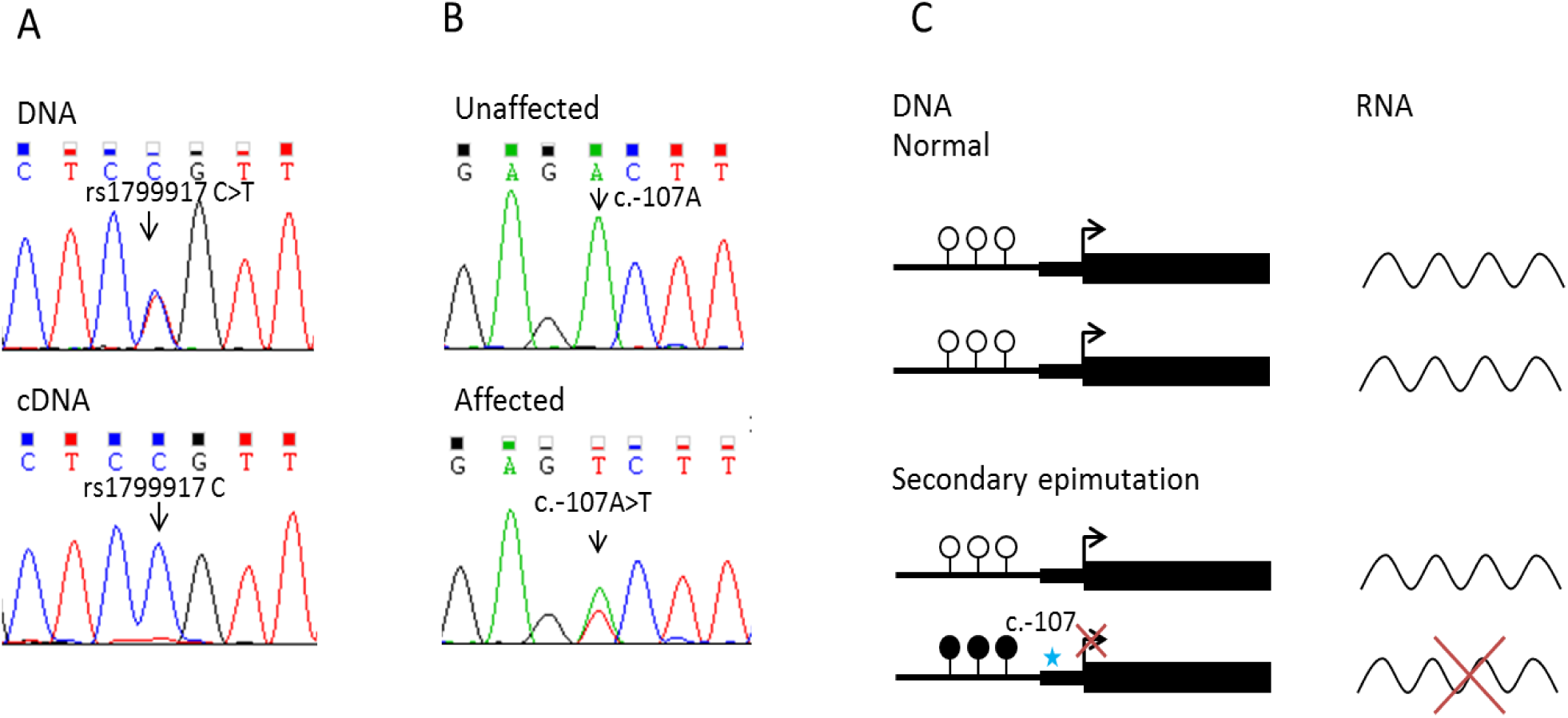
**A.** Representative Sanger sequencing traces demonstrating allelic loss of expression of rs1799917 C>T in exon 11 of *BRCA1*. In the DNA trace, both the C and T nucleotides are present, whereas in the cDNA trace only the C nucleotide is present. **B.** Representative Sanger sequencing traces for novel heterozygous c.-107A>T variant, present in individual with methylated *BRCA1* promoter and absent in an individual with an unmethylated promoter. **C.** Schematic representation of normal pattern of gene expression and transcription (top) and abnormal gene expression and transcription, caused by a germline variant (c.-107), resulting in hypermethylation of the promoter (secondary epimutation) and silencing of one allele.

Sanger sequencing upstream of the *BRCA1* translation start site identified a novel heterozygous variant c.-107A>T (g.43125358A>T, NM_07294.3) in an affected woman with *BRCA1* promoter hypermethylation in each family (Figure 3B). This variant segregated with the hypermethylated *BRCA1* allele in all tested individuals in both families and was absent in individuals lacking the hypermethylated allele, confirming that it was in *cis* (Table 1, eTable 2). This variant was absent in gnomAD^21^, a database that includes whole exome and whole genome sequencing data on 123,136 and 15,496 individuals, respectively. The variant has not been reported in any individual with breast or ovarian cancer in disease-specific databases, including the BRCA Exchange (http://brcaexchange.org).

The two families (both non-consanguineous white British from North West England) were not knowingly related to each other. All individuals in the two families with promoter hypermethylation and the c.-107A>T variant carried the previously described B1 haplotype (eTable 4, 5)^20^. To identify any additional germline variants that may result in promoter hypermethylation, we undertook SNP arrays and whole genome sequencing. SNP array analysis of II-5, III-2, and III-5 (Family 1), and III-2 and III-3 (Family 2) did not identify any other rare or novel copy number variants. Whole genome sequencing analysis was restricted to a candidate region (chr17:42,044,295-44,215,483, *hg38*) 1Mb upstream and downstream of the *BRCA1* gene. Segregation analyses were performed to identify variants in a heterozygous state in the two unrelated affected individuals (III-5, Family 1 and III-2, Family2) and absent in the unaffected individual (II-2, Family 1). This restricted analysis identified 14 variants that were absent from both the gnomAD dataset and dbSNP. Two novel variants (one in intron 2, c.80+661_80+667delAAAAAAA (g. 43123349-43123356delAAAAAAA, NM_007294.3) (eMethods, eFigure 3) and the previously identified c.-107A>T variant) were determined to be within the genomic region for *BRCA1*. Three variants within the candidate interval were present within DNase I hypersensitivity sites characterized across 125 cell types. In combination, these analyses identified c.-107A>T as single novel candidate variant linked to hypermethylation of the promoter (eFigure 2).

## Discussion

Here, over 20 years after the initial report that pathogenic variants in *BRCA1* result in familial breast cancer,^22^ we demonstrate for the first time transgenerational epigenetic silencing of *BRCA1* in two families with early-onset breast and ovarian cancer. A constitutional epimutation describes an epigenetic change, for example promoter hypermethylation, that results in the transcriptional silencing of a gene that is normally active across a range of normal tissues and predisposes to disease. Sloane *et al*.^23^ set out four criteria to establish the presence of a constitutional epimutation, which are met in our two families, in that promoter hypermethylation is confined to one allele in normal tissues derived from the mesoderm (blood), hair follicles (ectoderm), and buccal mucosa (endoderm); the level (~50%) and presence of hypermethylation is demonstrated using at least two independent methods (pyrosequencing and clonal bisulfite sequencing); the methylated allele is transcriptionally silent and there is co-segregation of the methylated and transcriptionally silent allele with the phenotype.^23^

Inherited epigenetic variants have only rarely been described in familial cancer, notably in Lynch syndrome due to hypermethylation of the *MLH1* promoter^15^, or *MSH2* promoter.^16^ *MLH1* promoter hypermethylation has been reported both in the context of a *cis*-acting germline variant, c.-27C>A (secondary epimutation), and in the absence of any detectable genetic alteration (primary epimutation).^24^ In contrast *MSH2* promoter hypermethylation has always been associated with a *cis*-acting deletion encompassing the 3’ end of the adjacent *EPCAM* gene.^16,25^ Here, we identified a novel *BRCA1* exon 1 variant, c.-107A>T, in *cis* with the hypermethylated promoter, which segregated with the phenotype in both families.

In this family there is, as yet, no evidence to determine if male to female vertical transmission of the *BRCA1* promoter methylation results in a breast/ovarian cancer phenotype in the next generation. Future predictive testing of the at-risk daughters of male carriers in this family will be able to establish this. However, as there is a linked upstream variant (c.-107A>T), it is likely that transmission will result in promoter methylation and a phenotype.

The c.-107A>T *BRCA1* variant is found on an ancestral B1 haplotype^20^ in both families. Although the families are not known to be related to each other, this data indicates that the two families may share a common ancestry. It will be important to determine if this variant occurs in other affected individuals to establish if this variant has arisen more than once and if other non-coding variants can result in *BRCA1* promoter hypermethylation. The c.-107 nucleotide is not highly conserved through mammalian species and *in silico* tools are not informative in predicting its pathogenicity. Notably exon one is not normally sequenced in clinical *BRCA1* testing and so the c.-107A>T variant would not have been detected by routine testing. Even if it had been identified by sequence analysis, without the methylation studies it would be classified as a variant of unknown significance. Therefore, promoter methylation studies are merited to clarify the functional effect of all rare or novel 5’ variants in *BRCA1*. The specific mechanism by which the 5’ variant results in promoter hypermethylation is, as yet, unknown.

Importantly, the clinical presentation of the affected individuals in the two families is consistent with the phenotype in other families with *BRCA1* pathogenic variants and does not indicate any specific clinical features that would prioritize individuals with familial breast/ovarian cancer without coding *BRCA1* pathogenic variants for methylation analysis. Although based on only two families the penetrance of the hypermethylated *BRCA1* promoter is 71·4% in informative women. This is consistent with estimates of cumulative risks by age 80 years for females with pathogenic *BRCA1* variants of 75% for breast cancer.^5^ The two unaffected female variant carriers were born before 1940 when penetrance for *BRCA1* pathogenic variants was much lower.^26^ For the male relatives, as expected, there is no evidence of an elevated cancer risk.^27^ Variable (mosaic) levels of *BRCA1* promoter methylation are detected in normal somatic tissues from individuals carrying the 5’ variant ranging from 24% in hair in individual II-5 in Family 1 to 69% in buccal mucosa in individual III-2 in Family 2, both women have bilateral breast cancer. There is no correlation between these promoter methylation levels and the clinical phenotype, for example the variant carriers (II-4) in Family 2 has methylation levels >40% but does not have cancer at 85 years. We detected the epimutation in two of 49 families ascertained in North West England with a Manchester score of >34. This equates to this mechanism accounting for at least 1.25% of *BRCA1* pathogenic variants in our very high-risk familial breast/ovary cohort and increases sensitivity from 71·8% to at least 72·7% in families with a high likelihood of a *BRCA1/2* pathogenic variant. Therefore, this mechanism is more common in our population than deep intronic mutations.^7^ Further, in our familial breast/ovarian cancer cohort, next generation sequencing of a panel of genes associated with an increased risk of breast cancer increased the diagnostic yield for familial breast cancer by a similar amount, but only revealed variants in genes (*ATM/CHEK2*) with less clear actionability.^5,7^ The uplift achieved by methylation testing would argue that *BRCA1* promoter methylation testing is a valuable adjunct to sequence and copy number analysis for individuals with a strong family history of breast and/or ovarian cancer.

In summary, we identified two families with transgenerational allele specific promoter methylation of *BRCA1*, which is present in all three germ layers, resulting in transcriptional silencing of one allele. The promoter hypermethylation is linked to an *in cis* variant c.-107T>A. This novel mechanism may explain some of the missing heritability in familial breast/ovarian cancer families.

## Acknowledgements

Many thanks to the family members for their participation and co-operation. Funded by Prevent Breast Cancer (GA 12-006 and GA 15-002) and the Manchester NIHR Biomedical Research Centre (IS-BRC-1215-20007). DGE is an NIHR Senior Investigator (NF-SI-0513-10076). JE is funded by a Post Doctoral Research Fellowship from Health Education England Genomics Education Programme. The views expressed in this publication are those of the author(s) and not necessarily those of HEE GEP. All funding bodies had no role in the design and conduct of the study; collection, management, analysis, and interpretation of the data; preparation, review, or approval of the manuscript; and decision to submit the manuscript for publication.

DGE and EvV had full access to all the data in the study and take responsibility for the integrity of the data and the accuracy of the data analysis. All tested individuals gave consent to present their personal information in this manuscript.

With thanks to Jeanette Rothwell for sample collection and Natasha Leo for technical assistance. The authors would like to thank the Genome Aggregation Database (gnomAD) and the groups that provided exome and genome variant data to this resource. A full list of contributing groups can be found at http://gnomad.broadinstitute.org/about.

TA declares an honorarium from Illumina for speaking. The other authors have no disclosures or conflicts of interest to declare. DGE, EvV, MS, and WN designed the study. DGE and WN obtained funding. DGE, DE, and FL ascertained patients and collated all relevant clinical information. EvV, MS, HB, AW, GB, JE, TA, and JSL carried out the experiments. All authors contributed to the analysis of the data. DGE, EvV, MS, and WN drafted the manuscript. All authors commented and approved the final version.

